# Sendai Virus Infection Induces Expression of Novel RNAs in Human Cells

**DOI:** 10.1101/451369

**Authors:** Roli Mandhana, Curt M. Horvath

## Abstract

Innate antiviral immune responses are driven by virus-induced changes in host gene expression. While much research on antiviral effectors has focused on virus-inducible mRNAs, recent genome-wide analyses have identified hundreds of novel target sites for virus-inducible transcription factors and RNA polymerase. These sites are beyond the known antiviral gene repertoire and their contribution to innate immune responses is largely unknown. In this study, RNA-sequencing of mock-infected and Sendai virus-infected cells was performed to characterize the virus-inducible transcriptome and identify novel virus-inducible RNAs (nviRNAs). Virus-inducible transcription was observed throughout the genome resulting in expression of 1755 previously RefSeq-annotated RNAs and 1545 nviRNAs. The previously-annotated RNAs primarily consist of protein-coding mRNAs, including several well-known antiviral mRNAs that had low sequence conservation but were highly virus-inducible. The previously-unannotated nviRNAs were mostly noncoding RNAs with poor sequence conservation. Independent analyses of nviRNAs based on infection with Sendai virus, influenza virus, and herpes simplex virus 1, or direct stimulation with IFNα revealed a range of expression patterns in various human cell lines. These phylogenetic and expression analyses suggest that many of the nviRNAs share the high inducibility and low sequence conservation characteristic of well-known primary antiviral effectors and may represent dynamically evolving antiviral factors.

## Introduction

Virus infection of human cells initiates host cell signaling cascades that result in widespread changes in gene expression. Collectively, these changes produce robust antiviral responses that form the basis of innate and adaptive immunity^1,2^. Induction of the immune response depends on recognition of viral components by host pattern recognition receptors. Though RNA- and DNA-genome viruses are detected by distinct pattern recognition receptors^3,4^, the cellular response to most virus infections culminates in the activation of master transcription factors IRF3 and NF-κB.

The most widely studied virus-induced RNAs encode effector proteins that mediate virus interference and signal amplification^5^, including several types of interferon (IFN), antiviral cytokines that are secreted from the infected cells^6^. Type I IFN includes the single IFNβ, as well as a number IFNα subtypes that all bind to a common IFNα/β receptor^6^; Type II IFN includes the single IFNγ that binds to the distinct IFNγ receptor^6^; Type III IFN includes IFNλ members, also known as IL-28 and IL-29, that bind to a distinct IFNλ receptor^7^. These receptors activate downstream JAK-STAT signaling^8,9^ and the expression of hundreds of effectors, known as interferon stimulated genes (ISGs)^10-12^.

The rise in genome-wide sequencing studies has revealed many previously unrecognized RNA species including noncoding RNAs with diverse roles in cellular processes such as gene expression, post-transcriptional regulation, translation, cell cycle regulation, and immune responses^13-17^. Noncoding RNAs are also increasingly being linked to the cellular response to viruses. For example, BISPR is an IFNα-induced long noncoding RNA (lncRNA) that regulates the expression of tetherin, a cellular antiviral factor that blocks viral release^18,19^. NEAT1 is a virus-induced lncRNA that regulates the expression of antiviral genes such as IL-8^20^. In the case of lncRNA-ACOD1, expression promotes rather than interferes with virus replication by regulating cellular metabolism^21^. These examples underscore the importance of deep sequencing approaches to characterize the extent of noncoding RNA transcription following virus infections.

Our previous study in human cells showed that early responses to RNA-genome virus infections initiate genome-wide binding of IRF3 and NF-κB to diverse loci to recruit and activate RNA polymerase II, assemble transcriptional machinery, and induce transcription^22^. Many of these binding sites were novel virus-inducible sites in intergenic and intronic regions of the genome, suggesting the existence of novel virus-inducible RNAs^22^. The identification of these novel binding sites along with the recent increase in identification of individual novel RNAs with functions related to regulation of virus infection motivated genome-wide examination of virus-inducible RNAs. In this study, novel RNA-encoding loci were identified throughout the genome, but unlike most annotated RNAs, they were primarily expressed from intergenic regions. Analysis of newly-identified virus-induced RNAs, referred to here as nviRNAs^22^, reveals that they can be induced by RNA and DNA-genome viruses as well as by direct stimulation with the type I IFN, IFNα, in both general and cell-specific fashion. These findings expand the extent of virus-induced transcription and suggest that both coding and noncoding RNA expression are hallmarks of the cellular response to virus infections.

## Results

### RNA-sequencing and identification of differentially expressed genes

To fully appreciate the extent of virus-induced RNA transcription, human Namalwa B cells were infected with Sendai virus, an RNA virus that is a potent inducer of antiviral signaling^22^. Mock-infected cells and cells infected with 5 plaque forming units (pfu) of Sendai virus per cell for 6 hours were subjected to paired-end RNA sequencing. Approximately 200 million reads of 100 bp read length were obtained for each sample. On an average, 90% of total reads mapped to the human reference genome (GRCh37/hg19).

Enrichment of RNAs in mock-infected or virus-infected samples was analysed with the DESeq2 program, resulting in the identification of 6210 RNAs that were differentially expressed after virus infection at a false discovery rate (FDR) of 5% and fold change of at least 1.5. Among these RNAs 3300 were induced by virus infection, while 2910 were suppressed. The expression changes of representative groups of these RNAs were confirmed in independent samples by RT-qPCR using primers specific for virus-induced previously-annotated genes (Fig. 1a) and previously-unannotated RNAs (Fig. 1b), as well as for RNAs suppressed by virus infection (Fig. 1c). In all categories, changes in RNA abundance levels measured by RT-qPCR closely matched the expression changes measured in RNA-sequencing analysis.

**Figure 1.**
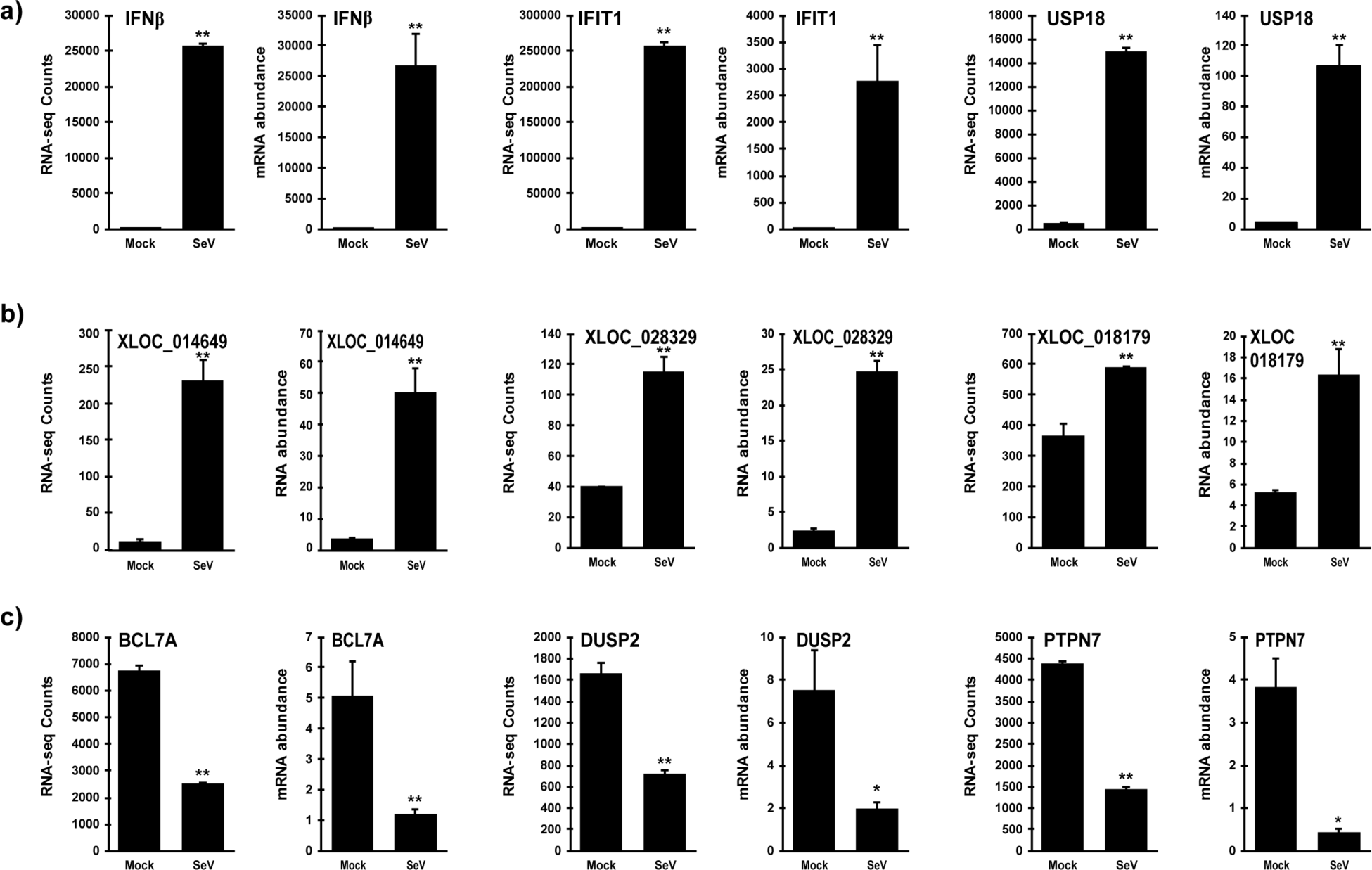
Validation of differential gene expression induced by Sendai virus infection. Namalwa cells were infected with Sendai virus for 6 hours and total RNA was analyzed by RNA-sequencing or by RT-qPCR in independent samples. For each indicated RNA, RNA-sequencing counts are plotted on the graph on the left and RNA abundance from RT-qPCR on the right. Expression of virus-induced a) previously-annotated genes and b) previously-unannotated RNAs and c) virus-suppressed genes was validated. RNA abundance data is representative of ≥ 2 replicate experiments and is shown normalized to GAPDH expression. Bars indicate average values of technical replicates (n=3) with error bars representing standard deviation. Statistical analysis was done using a two-tailed Student’s *t*-test for RT-qPCR measurements (* p-value < 0.05, ** p-value < 0.005)

The magnitude of differential expression was much less for the suppressed RNAs compared to the RNAs induced by virus infection. The expression of the most suppressed RNA, C10orf71, was suppressed only 50-fold by virus infection while the most induced RNA, IL-29, was induced almost 20,000-fold. More than 480 of the virus-induced RNAs are induced by more than 50-fold after virus infection. Due to this discrepancy in range of expression change, further emphasis was placed on analysing the virus-induced RNAs.

Of the 3300 virus-induced RNAs, 1755 have an existing RefSeq annotation^23^. The most highly induced of these annotated RNAs (Table 1) code for proteins known to play key roles in the cellular response to viruses, including type I IFNs^6^ (IFNβ, IFNα8, and IFNα13), type III IFNs^7^ (IL-29, IL-28α and IL28β), chemokines^24^ (CXCL10 and CXCL11), the antiviral mediator, OASL^25^, and the antiviral E3 ligase, HERC5^26,27^. In addition, 115 of the 389 previously identified type I ISGs compiled in a recent study^28^ were found to be induced by at least 1.5 fold.

**Table 1.** The top 10 highly virus-induced previously-annotated RNAs.

| Gene | Log<sub>2</sub> fold change | FDR | PhastCons mean score |
| --- | --- | --- | --- |
| IL29 | 12.21 ± 0.41 | 3.34E-191 | 0.187 |
| IFNβ | 12.05 ± 0.37 | 9.06E-229 | 0.209 |
| IL28β | 11.77 ± 0.40 | 1.33E-186 | 0.103 |
| CXCL11 | 11.66 ± 0.34 | 7.83E-250 | 0.209 |
| IL28α | 11.27 ± 0.42 | 3.42E-153 | 0.275 |
| OASL | 10.80 ± 0.19 | 0.00E+00 | 0.338 |
| CXCL10 | 10.15 ± 0.13 | 1.39E-168 | 0.238 |
| IFNα13 | 9.89 ± 0.45 | 0.00E+00 | 0.072 |
| IFNα8 | 9.73 ± 0.45 | 1.17E-106 | 0.094 |
| HERC5 | 9.72 ± 0.38 | 4.91E-102 | 0.495 |

### Gene enrichment analysis

To further characterize these induced RNAs, cellular pathways induced by virus infection were identified using DAVID Bioinformatics Resources^29,30^, a functional annotation tool that determines statistically significant gene enrichment in biological processes from a given gene list. The previously-annotated virus-induced RNAs mapped to 1608 DAVID gene IDs and were significantly enriched in more than 400 Gene Ontology (GO) biological processes and 40 Kyoto Encyclopedia of Genes and Genomes (KEGG) pathways.

As expected, the top 10 most enriched GO processes describe cellular functions that enable the cell to respond to virus infection, including type I IFN and cytokine signaling (Fig. 2a). The top 10 most enriched KEGG pathways also describe pathways in response to various RNA viruses such as influenza A virus, measles virus, and hepatitis C virus, a DNA virus, herpes simplex virus, and several pattern recognition receptor pathways including toll-like receptor signaling, RIG-I-like receptor signaling, and cytosolic-DNA sensing (Fig. 2b).

**Figure 2.**
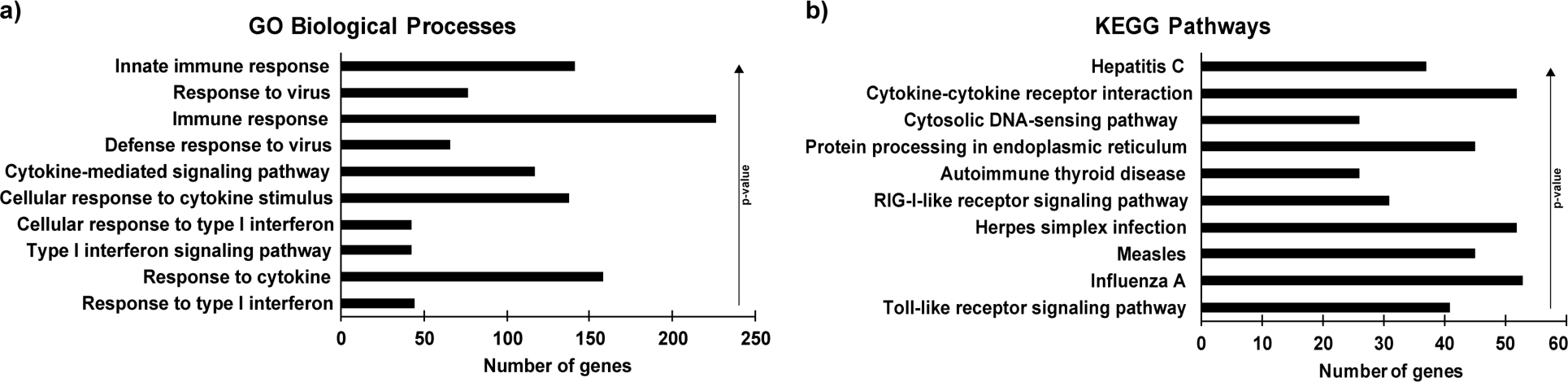
Graphical representation of functional enrichment analysis of Sendai virus-induced RNAs. Sendai virus-induced previously-annotated RNAs were analysed using DAVID to determine enriched a) GO biological processes and b) KEGG pathways. The top 10 most significant terms in each analysis are shown here in rank order by p-value. Bars represent number of virus-induced RNAs mapping to each term.

To better understand the extent of biological processes that are induced after virus infection, clustering analysis was carried out on all enriched GO biological process terms. The enriched GO biological processes formed 27 clusters with an enrichment score above 3, indicating that a majority of the processes within the cluster were statistically significant (Supplementary Table S1). The top five most enriched clusters relate to type I IFN signaling, cytokine signaling, type II IFN signaling and the unfolded protein response. The IFN and cytokine signaling pathways are known to induce antiviral factors, and the unfolded protein response is a cellular stress response induced by viral protein translation in the host cell. These analyses show that the previously-annotated virus-induced RNAs include many factors with known roles in the cellular response to viruses and that the experimental system used in this study is capable of uncovering proteins and pathways that are known to be important in the response to viruses.

### Genomic distribution of annotated RNAs and nviRNAs

Among the 3300 virus-induced RNAs there were 1755 previously RefSeq-annotated RNAs and 1545 novel virus-inducible RNAs (nviRNAs) that were not previously RefSeq-annotated. The previously-annotated RNAs and the identified nviRNAs were found to be widely distributed among all the chromosomes (Fig. 3a), but the nviRNAs differed from the known RNAs in their distribution among specific genomic features (Fig. 3b). Examination of six genomic feature categories, including: promoters, transcriptional termination sites, coding exons, introns, intergenic regions and untranslated regions (UTRs; includes 5’ and 3’ UTRs), revealed the majority of annotated RNAs mapped to coding exons (87.6%; Fig. 3b) with fewer than 10% mapping to introns and intergenic regions, indicating that the annotations largely reflect mRNA-encoding genes. In contrast, only 1.7% of the nviRNAs mapped to coding exons while more than 90% mapped to introns or intergenic regions. The finding that most nviRNAs are transcribed from intronic and intergenic regions is consistent with previous work showing abundant virus-induced RNA polymerase II binding and elongation throughout intronic and intergenic regions^22^.

**Figure 3.**
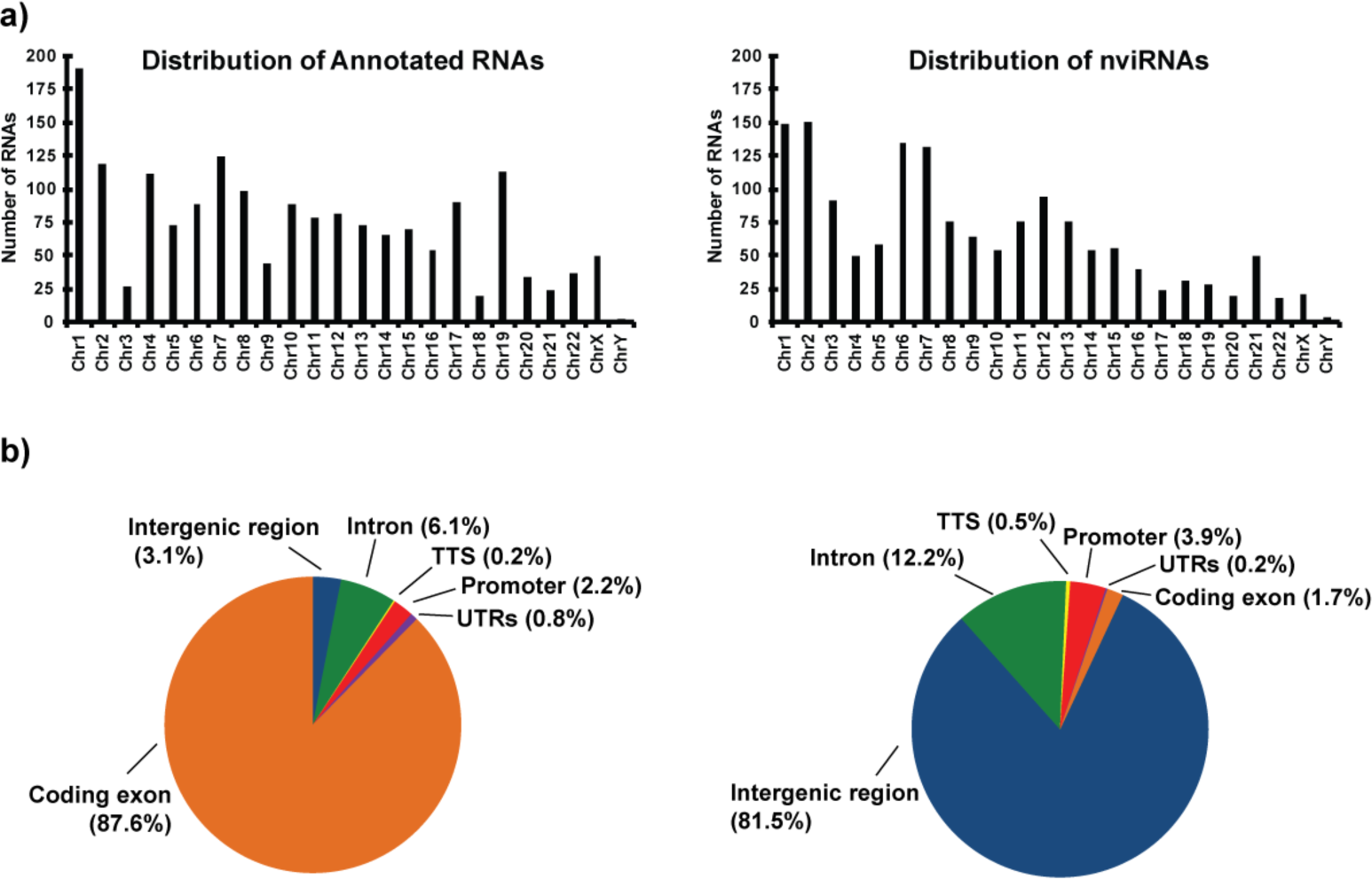
Comparison of genomic distribution of Sendai virus-induced previously-annotated RNAs and nviRNAs. a) Bar graphs represent the number of virus-induced previously-annotated RNAs (left) and nviRNAs (right) identified on each chromosome. b) Pie charts illustrate genomic feature distribution of previously RefSeq-annotated RNAs (left) and nviRNAs (right). RNAs are mapped to one of six annotation categories: promoters, transcriptional termination sites (TTS), exons, intergenic regions, introns, and untranslated regions (UTRs; includes 5’ and 3’ UTRs), with the percentage of sites corresponding to each category displayed in parentheses near the label.

### Protein coding and conservation analysis of annotated RNAs and nviRNAs

To compare the protein coding potential of the virus-induced RNAs, a PhyloCSF score was calculated for each RNA. PhyloCSF is a method that uses a multi-species sequence alignment to calculate a score reflecting the likelihood that an open reading frame (ORF) encodes a protein^31^. Analysis of previously characterized RNAs has shown that known noncoding RNAs generally score below 50 while protein-coding genes score above 50^32,33^.

There were 10 previously-annotated RNAs and 20 nviRNAs that did not have an ORF and were excluded from this analysis. The remaining 3270 RNAs were analysed and plotted in Fig. 4. Results indicate that the majority of the known RNAs (77.6%, Fig. 4a) were predicted to be protein coding mRNAs, while only 4.2% of the nviRNAs were predicted to encode a protein (Fig. 4a). Similar results were seen when limiting to RNAs induced at least 5-fold by virus infection (Supplementary Fig. S1). Of the previously-annotated RNAs, 69.6% were predicted to encode a protein while only 2.2% of the nviRNAs were likely to be protein coding. However, for both the previously-annotated RNAs and the nviRNAs, a larger proportion of the RNAs mapping to exonic genomic features had a high PhyloCSF score (Fig. 4b-c). For the annotated RNAs, 87% of the RNAs mapping to exons had a PhyloCSF score greater than 50 (Fig. 4b). Similarly, 52% of the nviRNAs mapping to exons had a PhyloCSF score above 50 (Fig. 4c). These results indicate that the nviRNAs are mostly virus-inducible noncoding RNAs, but there are 64 nviRNAs that were identified to have protein-coding potential.

**Figure 4.**
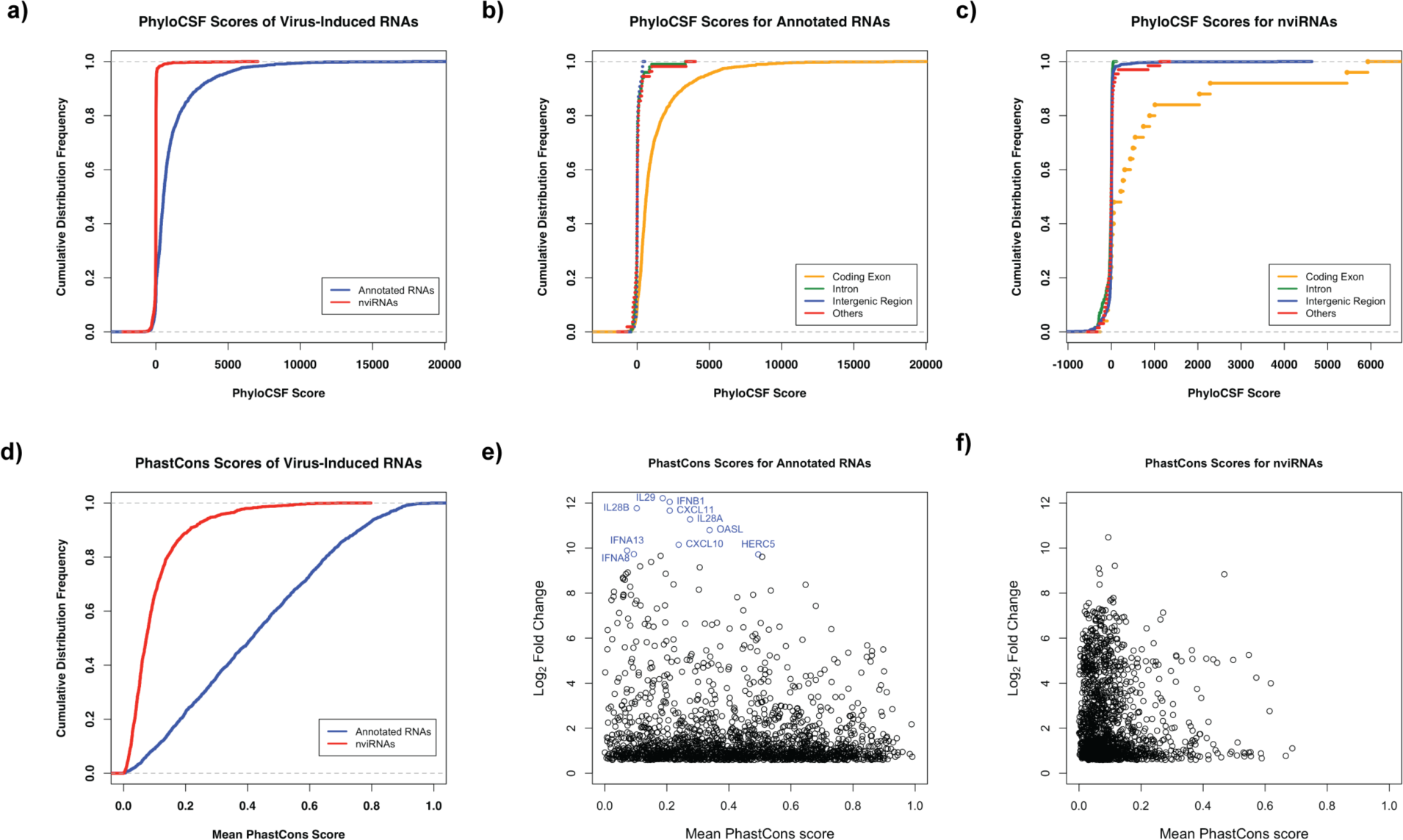
Comparison of protein coding potential and vertebrate sequence conservation of Sendai virus-induced previously-annotated RNAs and nviRNAs. a) The cumulative distribution frequency of PhyloCSF scores for previously-annotated RNAs (blue) and nviRNAs (red). b) The cumulative distribution frequency for RNAs mapping to the different genomic feature annotations for previously RefSeq-annotated RNAs and c) nviRNAs. The different genomic feature annotations include: coding exons (orange), intergenic regions (blue), introns (green), and promoters, transcriptional termination sites, and UTRs (all grouped together; red). d) The cumulative distribution frequency of the mean PhastCons score for the previously-annotated RNAs (blue) and nviRNAs (red). The mean PhastCons score for each RNA is plotted against the log_2_ fold change in expression after Sendai virus infection for e) previously-annotated RNAs and f) nviRNAs. The 10 most highly-induced previously-annotated RNAs are labeled in blue in e).

Though the nviRNAs are mostly noncoding RNAs, they may nevertheless have important functions in the antiviral system. Sequence conservation among vertebrates is frequently used to determine functional capacity of RNAs. However, many immune genes have undergone positive, diversifying selection and are therefore poorly conserved. Lower sequence conservation in virus-inducible genes may be indicative of an important antiviral role.

Sequence conservation among vertebrates was calculated by PhastCons, which uses a multiple-species alignment to calculate a score for each nucleotide corresponding to the probability that it is within a conserved element ^34^. To determine average sequence conservation across vertebrates, the mean PhastCons score for each RNA was calculated. Overall, the previously-annotated RNAs had much higher sequence conservation than the nviRNAs (Fig. 4d). While only 50% of the previously-annotated RNAs had a score of 0.413 or less, 98% of the nviRNAs had a score of 0.410 or less, indicating that their sequences were much less conserved than the annotated RNAs. The maximum mean PhastCons score for a nviRNA was 0.687, whereas there were 24 previously-annotated RNAs with scores of 0.9 or higher, suggesting that the nviRNAs on average represent a novel evolving group of RNAs.

Comparison of the PhastCons score with virus-inducibility revealed that the most highly induced previously-annotated RNAs were not highly conserved among vertebrates, in accordance with these genes having evolved under positive selection (Fig. 4e, blue circles). When limiting to RNAs induced at least 5-fold by virus infection, 50% of the previously-annotated RNAs had a score less than 0.343 (Supplementary Fig. S2). The top 10 most up-regulated annotated RNAs all had a mean PhastCons score under 0.5, with the top 5 having a score under 0.3 (Fig. 4e; Table 1). As there were examples of highly-induced nviRNAs that were similarly poorly conserved (Fig. 4f), it is anticipated that at least a fraction of these nviRNAs could have novel functions in the response to viruses.

### Inducibility of nviRNAs in other conditions

Most of the well-known antiviral genes, including type I IFNs, are broad-acting effectors that are widely induced by diverse viruses and in most cell types. However, there are some antiviral genes that have more restricted antiviral activity^28^.

To determine whether nviRNAs are activated in contexts other than Sendai virus infection, a subset of the nviRNAs were screened in Namalwa cells to determine whether their expression could also be induced by infection with two viruses identified through the KEGG pathways (Fig. 2): a distinct RNA-genome virus, influenza A virus (A/Udorn/72) and a DNA-genome virus, HSV-1. In addition, cells were subjected to direct stimulation with type I IFN (IFNα). Cells were either infected with 5 pfu/cell of virus for 10 hours or treated with 1000 U/mL IFNα for 6 hours prior to RNA isolation for gene expression analysis by RT-qPCR with gene-specific primers. The known virus-induced genes IFIT1^35^, IFNβ^6^, CSAG3 and USP18^36^ were used as controls. This analysis revealed great variation in the inducibility of the nviRNAs, not only in which stimuli induced them, but also in the level of induction (Fig. 5; Supplementary Table S2). Using a 2-fold increase in expression as a minimum for induction, the nviRNAs were classified into one of eight clusters based on their behavior (Fig. 5): (1) inducible by viruses and IFNα, (2) inducible by viruses only, (3) inducible by RNA-viruses and IFNα, (4) inducible by RNA viruses only, (5) inducible by Sendai virus and HSV-1 as well as IFNα, (6) inducible by both Sendai virus and HSV-1, (7) inducible by Sendai virus and IFNα, and (8) inducible by influenza A virus or HSV-1. The nviRNAs that were inducible by multiple viruses and IFNα may represent type I ISGs that are broad-acting novel antiviral effectors, while the nviRNAs induced by a subset of stimuli may represent restricted antiviral effectors.

**Figure 5.**
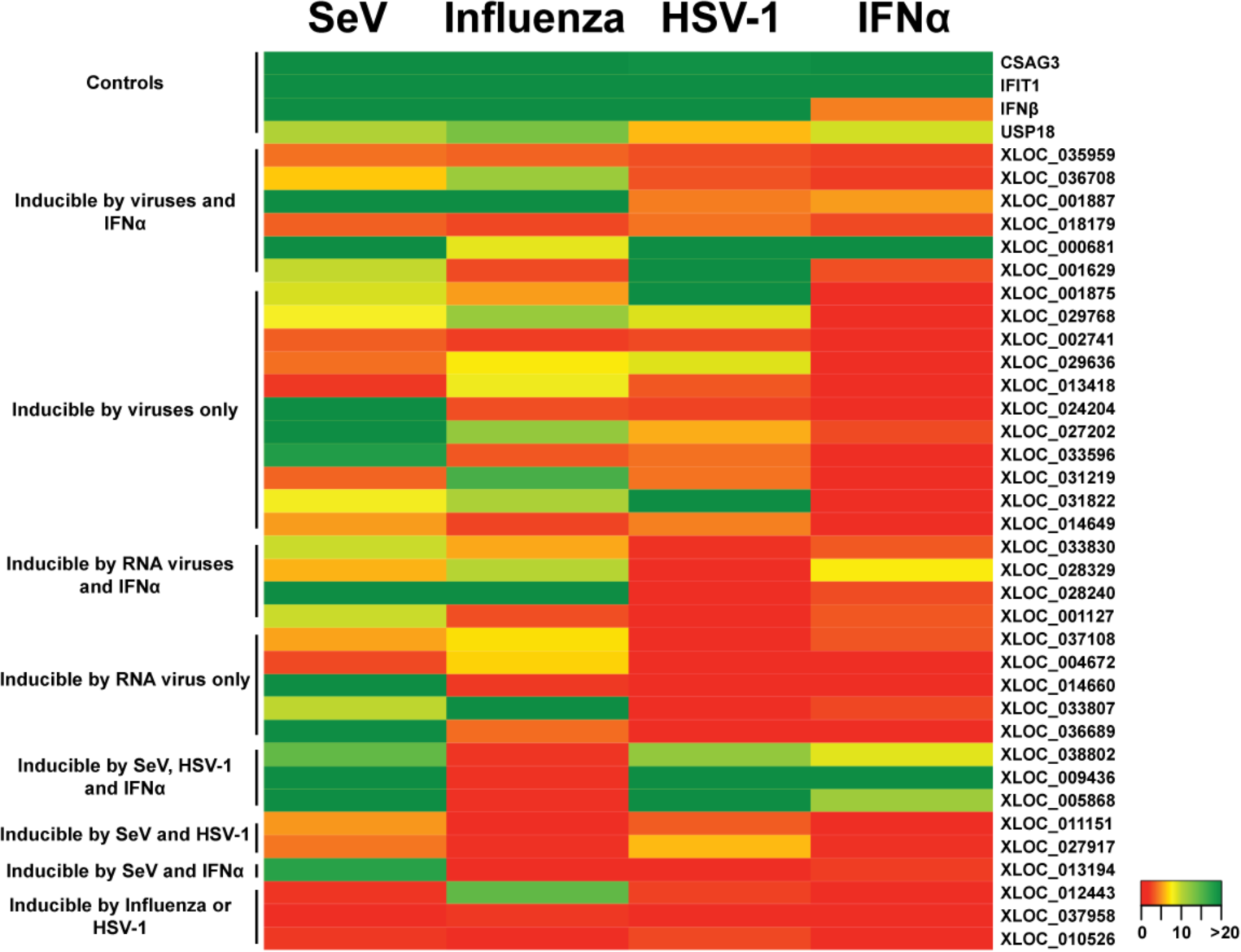
Classification of nviRNA expression in Namalwa cells from various stimuli. Total RNA from Namalwa cells infected with 5 pfu/cell for 10 hours of Sendai virus (SeV), influenza A virus, or HSV-1 or directly treated with 1000 U/mL of IFNα for 6 hours was analyzed by RT-qPCR. Heat map indicates expression of each RNA after infection or treatment with IFNα. Average values (n=3) of fold change are reported normalized to GAPDH expression.

To investigate the cell-specificity of these nviRNA transcriptional responses, the inducibility after virus infection and IFNα stimulation was also examined in several other human cell types. The epithelial 2fTGH cells and monocyte THP-1 cells were infected with 5 pfu/cell of Sendai virus (for 4 hours due to cytopathic effects) or influenza A virus (10 hours) or directly treated with IFNα (6 hours). THP-1 cells were also infected with 5 pfu/cell of HSV-1 (10 hours). In 2fTGH cells, transfection of the synthetic dsRNA, poly(I:C), an IFN activator that can robustly induce the expression of IFNs and ISGs, was included as a stimulus. The known virus-induced genes IFNβ^6^, IFIT1^35^, CSAG3 and USP18^36^ were expressed in both 2fTGH cells (Fig. 6a) and THP-1 cells (Fig. 7a) by various stimuli. The 6 nviRNAs identified as most likely to be broad-acting antiviral effectors in the Namalwa cells screen were also induced in both 2fTGH cells (Fig. 6b) and THP-1 cells (Fig. 7b). In 2fTGH cells, all the tested nviRNAs were induced by poly(I:C), though XLOC_035959 was induced less than 2-fold (Fig. 6b). Expression of the other 5 nviRNAs was also up-regulated more than 2-fold by influenza A virus. In THP-1 cells, influenza A virus robustly induced all 6 nviRNAs (Fig. 7b), with HSV-1 and IFNα also inducing half of the tested nviRNAs. The expression of the nviRNAs in the cluster only inducible by viruses in Namalwa cells was also tested in other cell types. There was greater variability in the inducibility of these nviRNAs. There were 6 nviRNAs induced in both 2fTGH cells (Fig. 6c) and THP-1 cells (Fig. 7c) by 10 hours of influenza A virus infection, while the other 5 nviRNAs showed more varied induction (Table 2).

**Figure 6.**
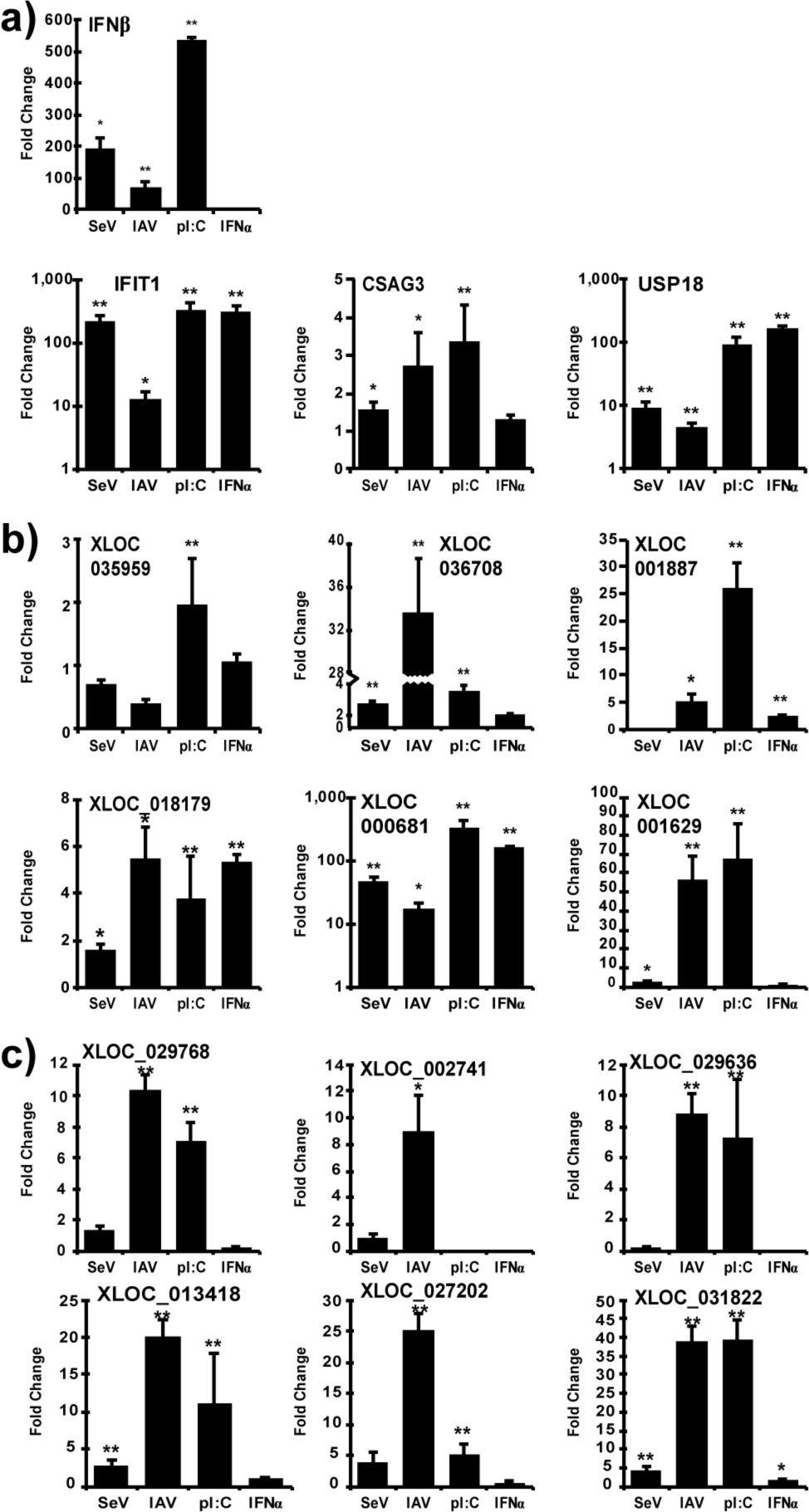
nviRNA expression in 2fTGH cells. Total RNA from 2fTGH cells infected with 5 pfu/cell Sendai virus (4 hours) or influenza A virus (IAV;10 hours), or transfected with synthetic dsRNA polyI:C (pI:C; 6 hours), or directly treated with IFNα (6 hours) was analyzed by RT-qPCR. Gene-specific primers were used for a) control genes, b) nviRNAs inducible by viruses and IFNα in Namalwa cells and c) nviRNAs only inducible by viruses in Namalwa cells. Data is representative of ≥2 replicate experiments and is shown normalized to GAPDH expression. Bars indicate average values of technical replicates (n=3) with error bars representing standard deviation. Statistical analysis was done using a two-tailed Student’s *t*-test (* p-value < 0.05, ** p-value < 0.005).

**Figure 7.**
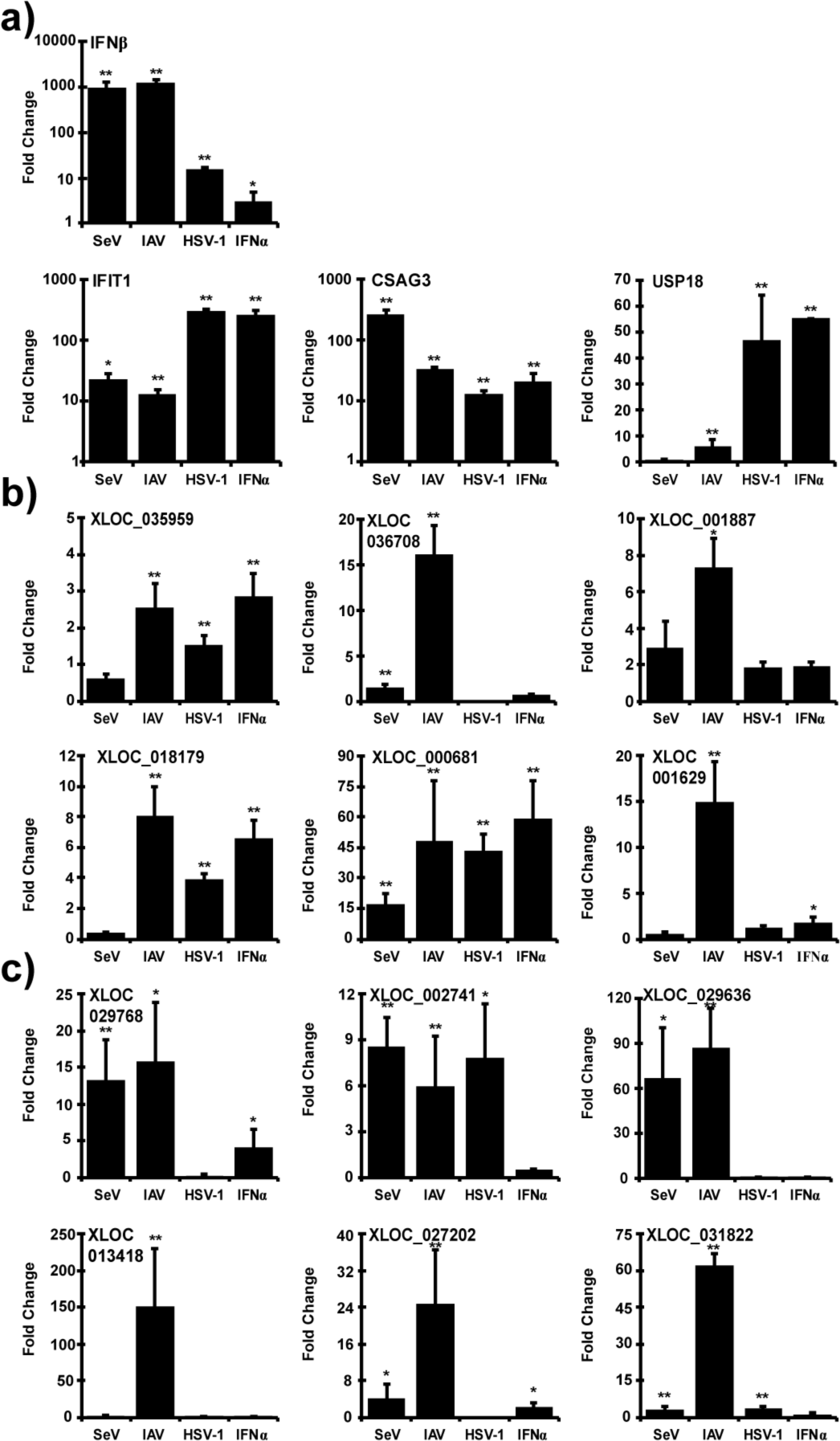
nviRNA expression in THP-1 cells. Total RNA from THP-1 cells infected with 5 pfu/cell Sendai virus (4 hours), influenza A virus (IAV; 10 hours) or HSV-1 (10 hours) or directly treated with IFNα (6 hours) was analyzed by RT-qPCR. Gene-specific primers were used for a) control genes, b) nviRNAs inducible by viruses and IFNα in Namalwa cells and c) nviRNAs only inducible by viruses in Namalwa cells. Data is representative of ≥2 replicate experiments and is shown normalized to GAPDH expression. Bars indicate average values of technical replicates (n=3) with error bars representing standard deviation. Statistical analysis was done using a two-tailed Student’s *t*-test (* p-value < 0.05, ** p-value < 0.005).

**Table 2.** Summary of nviRNA expression in various cell types.

|  | 2fTGH |  |  |  | HeLa |  |  |  |  | A549 |  |  |  | THP-1 |  |  |  |
| --- | --- | --- | --- | --- | --- | --- | --- | --- | --- | --- | --- | --- | --- | --- | --- | --- | --- |
| RNA | IFNα | SeV | IAV | pIC | IFNα | SeV | IAV | pIC | HSV-1 | IFNα | SeV | IAV | pIC | IFNα | SeV | IAV | HSV-1 |
| CSAG3 | - | - | + | + | + | + | + | + | + | + | + | + | + | + | + | + | + |
| IFIT1 | + | + | + | + | + | + | + | + | + | + | + | + | + | + | + | + | + |
| IFNβ | - | + | + | + | + | + | + | + | + | + | + | + | + | + | + | + | + |
| USP18 | + | + | + | + | + | + | - | + | + | + | + | + | + | + | - | + | + |
| XLOC_035959 | - | - | - | + | + | - | - | + | - | + | - | + | + | + | - | + | - |
| XLOC_036708 | - | + | + | + | - | - | + | + | + | - | - | + | + | - | - | + | - |
| XLOC_001887 | + | - | + | + | - | - | - | - | - | - | - | + | + | - | - | + | - |
| XLOC_018179 | + | + | + | + | - | - | - | + | - | + | - | + | + | + | - | + | + |
| XLOC_000681 | + | + | + | + | + | + | + | + | + | + | + | + | + | + | + | + | + |
| XLOC_001629 | - | + | + | + | + | - | + | + | + | - | + | + | + | - | - | + | - |
| XLOC_001875 | - | - | - | - | - | - | - | - | - | - | - | - | - | - | - | - | - |
| XLOC_029768 | - | - | + | + | + | - | + | + | + | - | - | - | - | + | + | + | - |
| XLOC_002741 | - | - | + | - | + | - | + | + | - | - | - | + | + | - | + | + | + |
| XLOC_029636 | - | - | + | + | - | - | - | + | - | - | + | + | + | - | + | + | - |
| XLOC_013418 | - | + | + | + | - | - | + | + | + | - | - | + | + | - | - | + | - |
| XLOC_024204 | - | - | - | + | - | - | - | - | - | - | - | - | - | - | - | - | - |
| XLOC_027202 | - | - | + | + | + | + | + | + | - | - | + | + | + | + | + | + | - |
| XLOC_033596 | - | - | - | - | - | - | - | - | - | - | - | - | - | - | + | + | + |
| XLOC_031219 | - | - | + | + | - | - | - | + | - | - | - | + | + | - | - | - | - |
| XLOC_031822 | + | + | + | + | + | - | + | + | + | + | + | + | + | - | + | + | + |
| XLOC_014649 | - | - | - | + | - | - | - | + | - | - | - | - | - | - | - | - | - |

The expression level and induction of each nviRNA varied greatly depending on the inducing stimulus, likely due to differences in the activation kinetics or intensity of each stimulus. Of the 12 nviRNAs expressed in both 2fTGH cells and THP-1 cells, 10 nviRNAs were also expressed in the epithelial HeLa and A549 cells (Table 2, Supplementary Table S3). The expression of these nviRNAs by at least 1 stimulus, and in most cases more than 1 stimulus, in multiple cell types demonstrates that these nviRNAs are not cell-specific and are candidates for antiviral function.

## Discussion

In this study, RNA-sequencing of Namalwa cells mock-infected or infected with Sendai virus identified 1755 previously RefSeq-annotated RNAs and 1545 novel virus-inducible RNAs (nviRNAs). Analysis of these virus-inducible RNAs revealed insights into the previously unrecognized depth and breadth of the virus-inducible transcriptome, and identified nviRNAs as possible targets of positive selection in the cellular response to viruses.

This study was motivated in part by the identification of novel virus-inducible binding sites for the antiviral transcription factors IRF3 and NF-κB, and RNA polymerase II^22^. Based on the RNA polymerase II binding sites, 1450 nviRNAs were predicted to be expressed^22^. Using the same cell line and virus for infection, 501 of these predicted nviRNAs were partially matched to RNAs identified in RNA-sequencing analysis. Factors that may contribute to this apparent disparity include timing, abundance and methodology. Furthermore, most of these RNAs were excluded from further analysis due to their low expression levels. Nevertheless, since many noncoding RNAs have low expression levels^37^, the subset of functional nviRNAs may extend past the 1545 nviRNAs analysed in this study.

Though both previously-annotated RNAs and nviRNAs were distributed similarly across the chromosomes (Fig. 3a), there was a significant difference in the genomic features associated with these different loci. More than 80% of the previously-annotated RNAs were transcribed from exons, whereas more than 80% of the nviRNAs were transcribed from intergenic regions (Fig. 3b). Even when limiting to RNAs with a high PhyloCSF score, indicating a strong likelihood that the RNA encodes a protein, 96.4% of the previously-annotated RNAs and only 21.8% of the nviRNAs mapped to exons. This discrepancy highlights that current annotations of genomic features are based on identification of a relatively small number of RNAs and the labeling of most of the genome as intergenic “junk” DNA. The recent expansion in sequencing studies that identify novel RNAs indicates a need for a corresponding refinement in annotations of genomic features.

The protein coding potential and functionality of the virus-inducible RNAs was assessed using PhyloCSF and PhastCons respectively. A majority of the previously RefSeq-annotated RNAs were identified as protein coding mRNAs, while very few of the nviRNAs were found likely to encode a protein (Fig. 4). The mean PhastCons score of each RNA was used to determine its average conservation. Though the previously-annotated RNAs vary in their mean PhastCons score, the most highly inducible, well-known RNAs that encode antiviral factors had a PhastCons score below 0.5 (Fig. 4e; Table 1), indicating low conservation may be suggestive of positive selection and overall importance to the cellular response to viruses.

While most genes have undergone negative selection to be conserved through evolution, many immune genes have been positively selected for diversity^38,39^. Virus evolution to overcome host antiviral immunity creates a potent selective force for diversification of key antiviral effectors. The cyclical virus-host interplay has resulted in immune genes being among the more recently evolved and more diversified genes in the human genome^40^. The poor sequence conservation of the highly induced nviRNAs may indicate that they are previously unidentified antiviral factors that evolved under positive selection.

In further support of this concept of nviRNA functionality, expression of a subset of nviRNAs was analysed and found to be induced, albeit variably, by different RNA and DNA viruses, and antiviral stimulation with poly(I:C) and IFNα. Differences in both expression and temporal regulation were observed among the nviRNAs tested due to specific virus or treatment, as well as in the cellular host cell line. It is reasonable to speculate that at least some of the nviRNAs, particularly the 10 expressed in 5 different cell lines tested (Fig. 5, Table 2), could represent novel antiviral factors with unique roles in the cellular innate immune response to viruses. This expression analysis does not assign any specific function to these novel RNAs, but their identification and classification is the first step in understanding their roles in the cellular response to virus infections. A loss of function analysis should be conducted for the nviRNAs that were classified as inducible in multiple cell lines to determine their function.

This study focused on identifying nviRNAs with potential functions in the cellular response to viruses; however, it is worthwhile to also examine potential novel antiviral factors among the previously RefSeq-annotated RNAs. Many of these RNAs encode known antiviral genes that are enriched in critical antiviral biological processes, but not all of the annotated RNAs have been assigned an antiviral function. Of the 1755 previously-annotated virus-induced RNAs, 328 were not assigned to any GO biological process, indicating that they are poorly characterized functionally. Additionally, there are many RNAs that despite being assigned a GO biological process, were not previously identified as virus-inducible RNAs. Combining these previously-annotated, virus-inducible RNAs with the nviRNAs identifies a large pool of potential antiviral responders that could have untapped value as antiviral diagnostic or therapeutic tools.

## Materials and Methods

### Cells, Viruses, and Treatments

Namalwa B cells (ATCC CRL-1432) and THP-1 cells (ATCC TIB-202) were cultured in RPMI 1640 Medium (Life Technologies, Thermo Fisher Scientific). HeLa cells (ATCC CCL-2), A549 cells (ATCC CCL-185) and 2fTGH cells (ECACC 12021508) were cultured in Dulbeco’s Modified Eagle Medium (Life Technologies, Thermo Fisher Scientific). For all cell lines medium was supplemented with 10% Cosmic calf serum (HyClone, GE Healthcare Life Sciences) and 1% Penicillin-Streptomycin (Gibco, Thermo Fisher Scientific). Sendai virus (Cantell strain) was grown in embryonated chicken eggs, and titers were determined on Vero cells. The A/Udorn/72 strain of influenza virus (gift of R. A. Lamb, Northwestern University) was propagated and titer was determined on MDCK cells. HSV-1 (F strain) (gift of G. A. Smith, Northwestern University) was propagated and titer was determined on Vero cells. Cells were infected at a multiplicity of infection (MOI) of 5 for indicated times. Virus infections were performed in serum-free medium (SFM), with 1% bovine serum albumin (BSA) supplemented for influenza virus. At 1 hour post-infection, the inoculation medium was replaced with medium containing 2% serum for the remainder of the infection. For poly(I:C) transfection, 2.5 μg/mL of low and high molecular weight poly(I:C) (Invivogen) was transfected for 6 hours using the Lipofectamine 2000 (Invitrogen, Thermo Fisher Scientific) transfection reagent. For direct treatment with IFNα, cells were treated with 1000 U/mL recombinant IFNα (Roche).

### Total RNA Extraction

Total RNA was extracted using the TRIzol Reagent (Invitrogen, Thermo Fisher Scientific) and treated with DNAse I (Invitrogen, Thermo Fisher Scientific). The RNA quantity was obtained using NanoDrop 2000 (Thermo Fisher Scientific). For RNA used in RNA sequencing, the integrity was examined by the Bioanalyser 2100 system (Agilent Technologies) to obtain RNA Integrity Number (RIN) of over 9.

### RNA sequencing and transcript assembly

Duplicate samples of mock-infected and Sendai virus-infected cells for 6 hours were used to prepare RNA libraries for sequencing on Illumina HiSeq2000 platform (Illumina) to generate 100 bp paired-end sequencing reads. Raw data was filtered to remove adapter sequences and low-quality reads. The remaining rRNA reads were removed by mapping to known human rRNA sequences. The clean, high-quality data was mapped to the human reference genome (GRCh37.p10/hg19)^41^ using TopHat^42^ (2.0.10). The mapped reads for each sample were independently assembled into annotated and novel transcripts using the Cufflinks^43^ (2.1.1) suite of programs.

### Bioinformatic analysis

To determine differential expression between mock-infected and Sendai virus-infected samples, read counts for each RNA were generated by HTSeq^44^ (0.6.0) for each sample and analysed by the DESeq2^45^ (1.2.10) program. Significance was calculated using the Wald test and a Benjamini-Hochberg False Discovery Rate cut-off of 5% was used to assess statistically significantly differential expression. The lowest quartile of RNAs based on expression were excluded from further analysis. RefSeq genomic feature distribution information for coding exons, introns, 3’ and 5’ untranslated regions (UTRs), promoters (−1kb), and transcription termination sites (+1 kb) was downloaded from the UCSC genome browser (http://genome.ucsc.edu/)^23^ and analysed using the BEDtools program^46^. The online functional annotation tool, DAVID^29,30^ (6.8), was used to conduct gene enrichment analysis. A list of gene symbols corresponding to differentially expressed RNAs was mapped to DAVID gene IDs to determine which Gene Ontology biological processes and KEGG pathways were enriched. A Benjamini-Hochberg False Discovery Rate cut-off of 5% was used to assess statistical significance. Clustering analysis of enriched GO biological processes was done using DAVID’s heuristic fuzzy multiple-linkage partitioning method^47^. An enrichment score cut-off of 3 was used to determine significance. Base-by-base PhastCons^34^ conservation score across 100 vertebrates were downloaded from the UCSC Genome Browser. For each RNA a mean score was calculated if there was a score available for at least 80% of the bases in the sequence. The PhyloCSF^31^ software was locally installed and used to determine the coding potential of the longest start-to-stop open reading frame in each RNA. The multiple-species alignments needed for this analysis were prepared on Galaxy web platform at usegalaxy.org^48^.

### cDNA synthesis and Quantitative PCR

Purified total RNA was random primed and reverse transcribed using SuperScript III Reverse Transcriptase (Invitrogen, Thermo Fisher Scientific). Gene expression was measured by quantitative real-time PCR (qPCR) and normalized to glyceraldehyde 3-phosphate dehydrogenase (GAPDH) to determine relative abundance by the 2^−⋄C^_T_ method or fold change over mock by the 2^−^⋄⋄CT method^49^. Primers are listed in Supplementary Table S4.

### Data Availability

RNA-sequencing data is available in the GEO database under study GSE115266.

## Acknowledgements

This work was supported by the Northwestern University NUSeq Core Facility, High Throughput Analysis Laboratory and Quest High Performance Computing Facility. We are grateful to Adam J. Hockenberry and members of the Horvath lab for critical comments and helpful suggestions, and to Dr. Christian Stehlik and Dr. Lucia Maria de Almeida for providing THP-1 cells. Supported by NIH grants AI073919, AI50707, and GM111652 to C.M.H. R.M. was supported by a pre-doctoral fellowship from the NIH Training Program in Viral Replication T32AI60523.

## Author Contributions

R.M. and C.M.H. designed the study and wrote the manuscript; R.M. conducted the experiments and analysis. All authors read and approved the final manuscript.

## Additional Information

### Accession codes

RNA-sequencing data is available in the GEO database under study GSE115266.

